# Predicting the conservation status of Europe’s Data Deficient sharks and rays

**DOI:** 10.1101/614776

**Authors:** Rachel H.L. Walls, Nicholas K. Dulvy

## Abstract

Shark and ray biodiversity is threatened primarily by overfishing and the globalisation of trade, and Europe has been one of the most documented heavily fished regions for a relatively long time. Yet, we have little idea of the conservation status of the hundreds of Data Deficient shark and ray species. It is important to derive some insight into the status of these species, both to understand global extinction rates and also to ensure that any threatened Data Deficient species are not overlooked in conservation planning. Here, we developed a biological and ecological trait model to predict the categorical conservation status of 26 Northeast Atlantic and 15 Mediterranean Sea Data Deficient sharks and rays. We first developed an explanatory model based on all species *evaluated* on the International Union for Conservation of Nature (IUCN) Red List of Threatened Species^TM^, using maximum body size, median depth (as a proxy for fisheries exposure), and reproductive mode, and then *predicted* the status of all Data Deficient species. Almost half of Northeast Atlantic (46%, *n*=12 of 26), and two-thirds of Mediterranean (67%, *n*=10 of 15) Data Deficient species are predicted to be in one of the three IUCN threatened categories. Northeast Atlantic Data Deficient species are *predicted* to be 1.2 times more threatened than *evaluated* species (38%, *n*=36 of 94), whereas threat levels in the Mediterranean Sea are relative for each (66%, *n*=38 of 58). This case study is intended for extrapolation to the global shark and ray dataset upon completion of the global IUCN Red List assessment. Trait-based, categorical prediction of conservation status is a cost-effective approach towards incorporating Data Deficient species into (i) estimates of lineage-wide extinction rates, (ii) revised protected species lists, and (iii) Red List Indices, thus preventing poorly known species from reaching extinction unnoticed.

## 1 INTRODUCTION

Despite a broadening of coverage of species and more intensive Red List assessment by the International Union for Conservation of Nature (IUCN) in the past decade, over one-sixth or around 13,465 species have been found to be Data Deficient (Bland et al., 2017). Data-deficiency is most prevalent in reptiles and amphibians, marine and freshwater organisms, invertebrates, and plants (Bland et al., 2012, 2014; Böhm et al., 2013; Callmander et al., 2005; Collen et al., 2012; Hoffmann et al., 2010). The IUCN classification means that there are insufficient data to make a more refined determination, hence Data Deficient species could range from actually being Least Concern or they could be threatened or even Extinct. Data-deficiency creates uncertainty in estimates of extinction rates, which is a key challenge to track progress towards the Convention on Biological Diversity’s (CBD) Aichi Target 12: to halt the loss of biodiversity by 2020 (CBD & UNEP, 2011). Clearly, a complete understanding of which species are threatened (Vulnerable, Endangered, or Critically Endangered) is an essential first step toward tracking and improving species’ status (Bland et al., 2014, 2015).

Data Deficient species are typically overlooked in conservation planning (Bland et al., 2014), with the implicit assumption that the biology and threatening processes of both Data Deficient and data-sufficient species are similar. To provide a first-approximation of the extinction rate of any taxon, the IUCN assumes Data Deficient species are equally as threatened as the data-sufficient species within a taxonomic group (Hoffmann et al., 2010). However, there are numerous reasons why the trait distribution and exposure to threatening processes might be different. For example, most recently discovered sharks have been found in the deep sea (Randhawa et al., 2015) and are relatively small-bodied, beyond the reach of most fisheries, hence those Data Deficient deepwater species may actually be Least Concern because they have refuge from the main threatening process of overfishing. Conversely, many recently resolved species complexes, such as devil rays, eagle rays, and skates may be highly exposed to fisheries and hence the newly described ‘Data Deficient’ species might already be highly threatened (Iglésias et al., 2010; White & Last, 2012).

There is a vast body of work on the correlates of population trajectories and extinction risk (Cardillo et al., 2005; McKinney, 1997; Owens & Bennett, 2000). Broadly, large body size, small geographic range, and ecological specialisation are the biological traits most often related to extinction risk, depending on their interaction with the appropriate threatening process (Owens & Bennett, 2000; Reynolds et al., 2005b). Only recently has this knowledge been used to predict extinction risk of Data Deficient species (Bland et al., 2015; Butchart & Bird, 2010; Dulvy et al., 2014; Jetz & Freckleton, 2015). Trait-based predictions of IUCN conservation status use biological and ecological trait data to predict the most likely categorisations for Data Deficient species based on assessed species. The simplest approach is to make the binary prediction whether a Data Deficient species is Least Concern or threatened. This approach has been used with a high degree of accuracy for mammals, birds, sharks and rays (Bland et al., 2015; Butchart & Bird, 2010; Dulvy et al., 2014; Jetz & Freckleton, 2015). The most significant advance has been the development of ordinal (or categorical) regression which enables prediction of the actual IUCN Red List category, based on relevant biological and ecological traits (Luiz et al., 2016). A total of 50 of 163 groupers (family Epinephelinae) were Data Deficient, yet trait-based ordinal regression revealed a total of three species predicted to be Critically Endangered, five to be Endangered, and 12 to be Vulnerable (Luiz et al., 2016).

Sharks and rays represent the oldest evolutionary radiation of vertebrate Classes (Stein et al., 2018), with an incredibly broad range of life-histories, spanning all ocean basins, and down to great depths (Cortés, 2000; Dulvy et al., 2014; Dulvy & Forrest, 2010). This makes them ideal for trait-based predictive modelling, while their high levels of population-relevant data-deficiency present the opportunity to test categorical predictions on a highly Data Deficient group for the first time. Europe represents the first region to be reassessed as part of an ongoing global IUCN Red List reassessment of sharks and rays, as well as being one of the most relatively data-sufficient regions for the Class (Dulvy et al., 2016; Fernandes et al., 2017; Nieto et al., 2015).

Here, we use Europe’s sharks and rays to consider three questions: (1) which biological and ecological traits are driving extinction risk; (2) how does the proportion of *evaluated*-threatened species compare with *predicted*-to-be-threatened Data Deficient species; and (3) which are the most threatened Data Deficient sharks and rays? We used cumulative link mixed-effects modeling (CLMM) to evaluate the relationship between species’ trait data and conservation status, and eventually predict the conservation status of Europe’s Data Deficient sharks and rays. This CLMM approach maintains the hierarchy of the IUCN categories while preventing the loss of information inevitable from lumping categories together as threatened and non-threatened (Luiz et al., 2016). Model performance was evaluated using the Akaike Information Criterion (AIC) with small sample size correction (AIC_c_).

## 2 METHODS

First, we describe the IUCN Red List conservation assessment of European sharks and rays. Second, we describe the development of an explanatory trait-based model to explain conservation status. Third, we describe the prediction and cross-validation of the conservation status of Europe’s Data Deficient sharks and rays.

### 2.1 IUCN Red List assessment

The European Red List assessments spanned the Northeast Atlantic Ocean and the Mediterranean and Black Seas, including the territorial waters and Exclusive Economic Zones of all European countries in the Northeast and Eastern Central Atlantic Ocean, and the offshore Macronesian island territories belonging to Portugal and Spain (Dulvy et al., 2016; Fernandes et al., 2017; Nieto et al., 2015).

In total, 131 species were assessed at the regional level for Europe using the 2001 IUCN Red List Categories and Criteria, version 3.1 (IUCN, 2012b). We convened 54 experts, composed mainly of members of the IUCN Shark Specialist Group, and completed the 131 European assessments over 21 months, from 2013–15. This culminated in a one-week workshop, attended by fifteen IUCN Shark Specialist Group members, to finalise and review all assessments. The assessed species included 50 skates and rays (Order Rajiformes), 72 sharks (Order Carcharhiniformes, Hexanchiformes, Lamniformes, Squaliformes, Squatiniformes), and nine chimaeras (Order Chimaeriformes). Only breeding residents of Europe were included in the assessments, including ‘visitor’ species defined by the IUCN as “a taxon that does not reproduce within a region but regularly occurs within its boundaries either now or during some period of the last century” (IUCN, 2012a). The only visitors in Europe are currently the Smalltooth Sawfish (*Pristis pristis*, Linnaeus 1758), and Largetooth Sawfish (*Pristis pectinata*, Latham 1794). Vagrant species were not included in assessments, which by IUCN definition are “a taxon that is currently found only occasionally within the boundaries of a region”. Vagrant species previously listed in Europe were listed as Not Applicable, and discounted from the following analyses (e.g., the Nurse Shark, *Ginglymostoma cirratum*, Bonnaterre 1788; Nieto et al., 2015). The IUCN Red List categories considered in this assessment are Least Concern, Near Threatened, Vulnerable, Endangered, Critically Endangered, and Data Deficient, as there are no sharks or rays known to be Regionally Extinct from the entire European region at present. This ordering represents lowest to highest extinction risk, with the exception of Data Deficient, which could include species that are both low and high risk.

### 2.2 Developing an explanatory trait-based model for conservation status

We considered three biological and ecological traits: maximum body size, median depth, and reproductive mode (Dulvy & Forrest, 2010; Dulvy & Reynolds, 2002; Field et al., 2009; Rigby & Simpfendorfer, 2015). Large maximum body size has been related to a greater likelihood of decline and extinction risk due to higher catchability and slower population growth rates in fishes, and other vertebrates (Dillingham et al., 2016; Field et al., 2009; Pardo et al., 2016). Deeper depth ranges are associated with refuge from fishing activity, and hence, lower extinction risk (Dulvy et al., 2014; Luiz et al., 2016). Overfishing is the greatest threat to sharks and rays and occurs predominantly down to 400 m deep and exceptionally down to greater depths (Bailey et al., 2009). Egg-laying (oviparous) species tend to be more fecund than live-bearing (viviparous) species and hence may have greater maximum population growth rates, greater variance in reproductive output, and hence scope for density-dependent compensation and lower sensitivity to fishing mortality for adults (Dulvy & Forrest, 2010; Forrest et al., 2008).

There are inherent differences in biogeography, fisheries, and fisheries management between Europe’s major sub-regions, the Northeast Atlantic Ocean and Mediterranean Sea, which warranted building models separately for each. There are 120 Northeast Atlantic sharks and rays and 73 Mediterranean species, so lumping the two together as a Europe-wide status created a bias towards Northeast Atlantic status. The IUCN categories were scored as Least Concern = 0, Near Threatened = 1, Vulnerable = 2, Endangered = 3, and Critically Endangered = 4 (Butchart et al., 2007). For each sub-regional model, IUCN category was the response variable and maximum body size (cm, total length), median depth (m), reproductive mode (scored oviparous = 1 or viviparous = 0). Median depth was used as a proxy for minimum depth and depth range to account for exclusively shallow or deep species’ distributions, while also avoiding having two highly correlated fixed effects within a model. We also considered the interaction between size and depth as a fixed effect. The interaction between size and depth is important because large-bodied species are only associated with higher extinction risk if they exist within the reach of fisheries (Dulvy et al., 2014). Size and depth were centred and scaled by two standard deviations. Family was included as a random effect to account for phylogenetic covariation.

### 2.3 Predicting conservation status

Predictive accuracy of the explanatory model was evaluated using Area Under the Curve (AUC) from Receiver Operating Characteristic curves (Sing et al., 2005). The AUC measure only works for binary classification, so to test the predictive accuracy of each of the five categories individually we scored each of the five IUCN categories separately as one, against all four other categories scored as zero. We also grouped the threatened categories (Critically Endangered, Endangered, Vulnerable) with a score of one, and non-threatened (Near Threatened, Least Concern) scored as zero to determine the model accuracy for predicting threatened versus non-threatened species. Predictive power was tested using data-sufficient species by dropping species from the model to predict the conservation status and cross-validate against each known, assessed conservation status. Test sets were run, comprising all data-sufficient species with all species dropped one at a time. The model for each sub-region that was able to predict the correct IUCN status with the highest AUC predictive accuracy measure was then used to predict the categories of the actual Data Deficient species. The highest overall accuracy for a model was determined by calculating the mean across all five AUC values for each IUCN category. Finally, the IUCN categorisation for each Data Deficient species was classified using a 50% cut-off point. All analyses were conducted in R version 3.5.2 (R Core Team, 2018), models were fit using the clmm2 function from the ordinal package (Christensen, 2019), and performance was evaluated with the ROCR package, version 1.0-7 (Sing et al., 2005).

## 3 RESULTS

### 3.1 IUCN regional European Red List assessment

One-fifth of the 120 Northeast Atlantic (22%, *n*=26) and 73 Mediterranean Sea (21%, *n*=15) shark and ray species assessed in 2015 are listed as Data Deficient (Figure 1, Table 1). Most species are assessed as Least Concern (38%) in the Northeast Atlantic, whereas the majority of species are Critically Endangered (27%) in the Mediterranean Sea (Figure 1). Specifically, of the data-sufficient Northeast Atlantic assessments, 38% (*n*=46) of species are Least Concern, 10% (12) Near Threatened, 8% (9) Vulnerable, 13% (15) Endangered, and 10% (12) Critically Endangered (Figure 1, Table 1). In the Mediterranean Sea, only 16% (12) species are Least Concern, 11% (8) Near Threatened, 10% (7) Vulnerable, 15% (11) Endangered, and 27% (20) Critically Endangered (Figure 1, Table 1). Sharks and rays are more threatened in Europe than the global average (17.4%, *n*=181; Table 1). Specifically, nearly one-third (30%, *n*=36) are threatened in the Northeast Atlantic and over half (52%, *n*=38) are threatened in the Mediterranean Sea. Rays are approximately as threatened as sharks in both regions, in the Northeast Atlantic 32% (*n*=14) of rays are threatened versus 33% (*n*=22) of sharks, and in the Mediterranean Sea 50% (*n*=16) of rays are threatened versus 55% (*n*=22) of sharks (Table 1).

**Figure 1.**
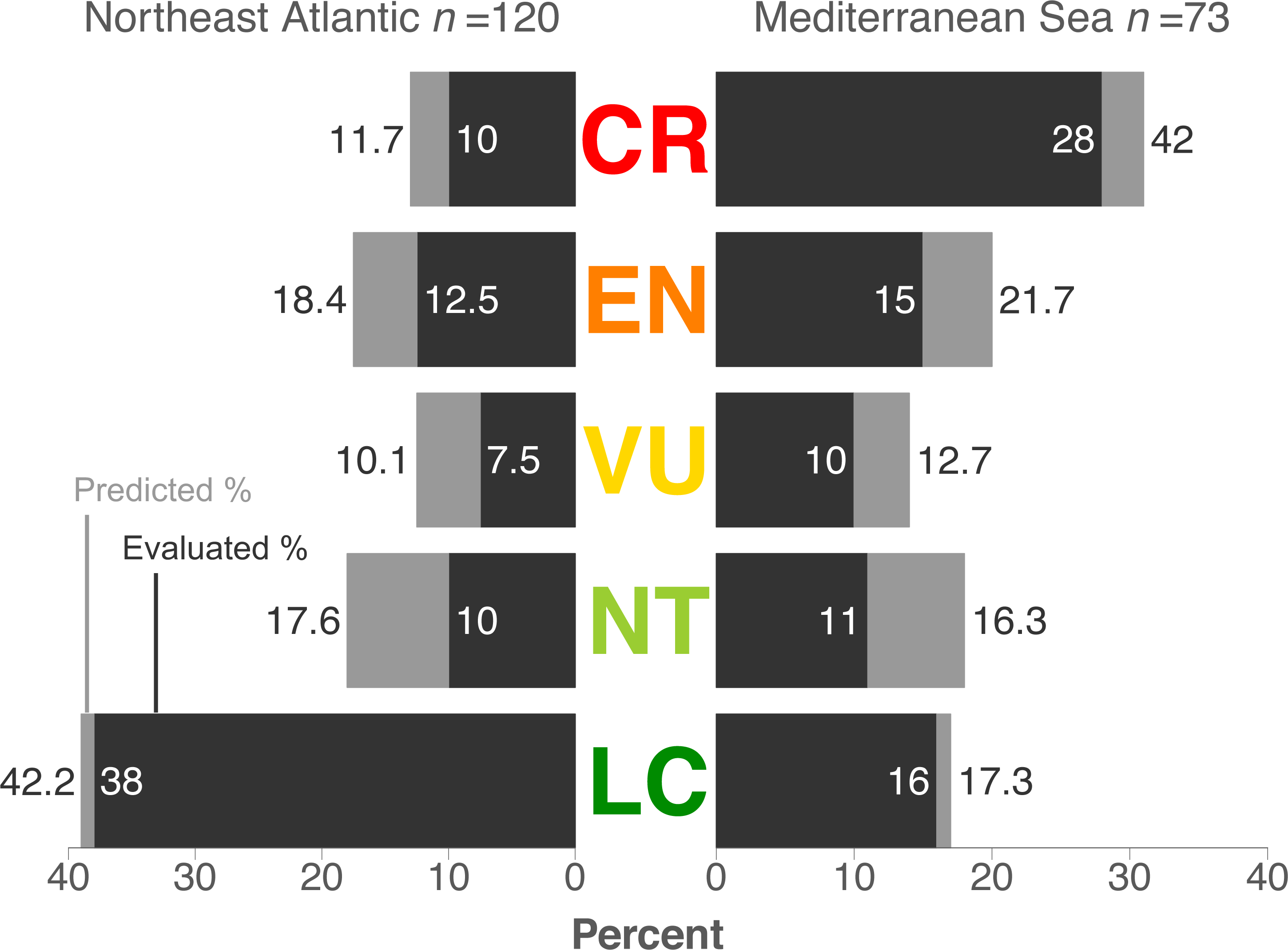
Percent *evaluated* and *predicted* IUCN categorisations of Europe’s sharks and rays. Dark bars represent the percentage of species officially evaluated on the IUCN Red List, while light bars represent the percentage of Data Deficient species predicted to be under each category as per the results of the present study. Of the 120 species in the Northeast Atlantic, 94 were *evaluated* and 26 were Data Deficient and *predicted* for. In the Mediterranean Sea, 58 of 73 species were *evaluated* and 15 were Data Deficient and *predicted* for. The IUCN categories from highest to lowest threat are: CR = Critically Endangered, EN = Endangered, VU = Vulnerable, NT = Near Threatened, and LC = Least Concern.

**Table 1.**
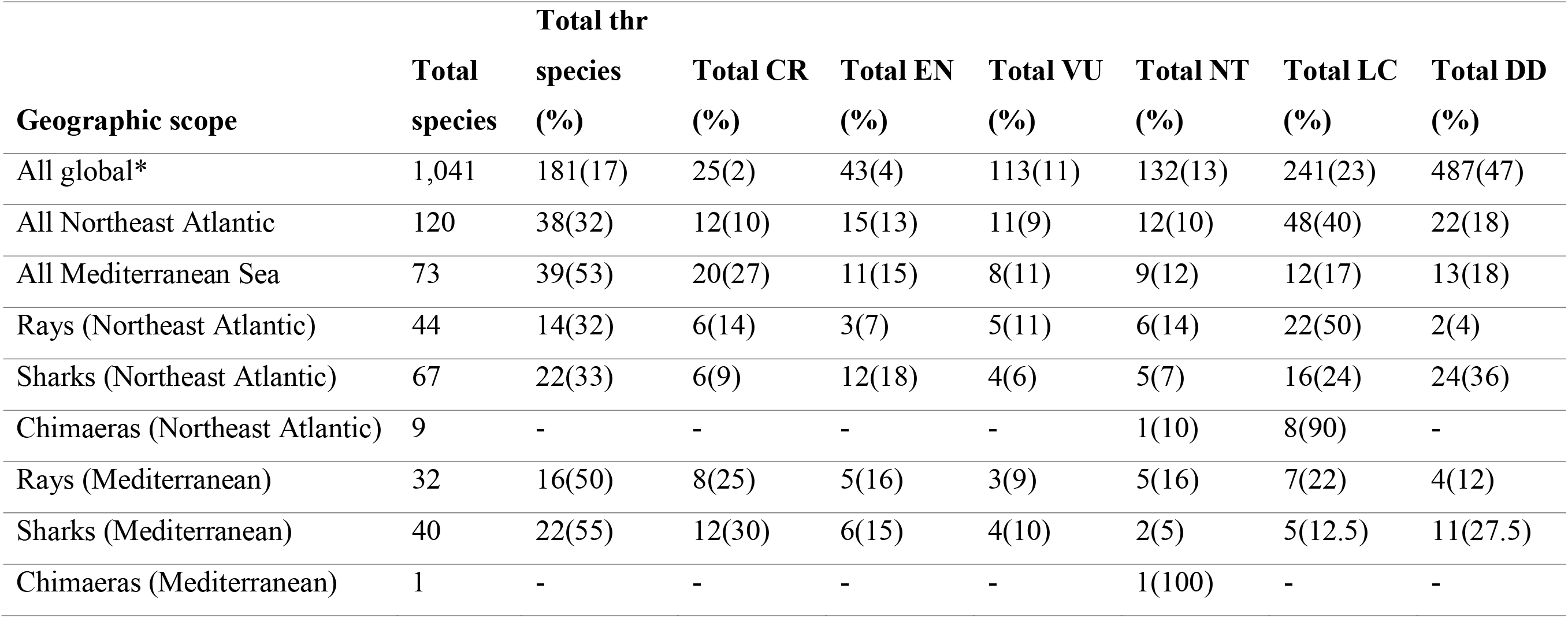
Global and European IUCN Red Listings of sharks and rays. Observed number and (percent) of global (2014), Northeast Atlantic (2015), and Mediterranean Sea (2015) sharks, rays (i.e. all rays and skates), and chimaeras in each IUCN Red List category. CR: Critically Endangered, EN: Endangered, VU: Vulnerable, thr: threatened (CR+EN+VU), NT: Near Threatened, LC: Least Concern, DD: Data Deficient (*Dulvy et al., 2014).

### 3.2 Biological and ecological predictors of conservation status

Large-bodied sharks and rays are more likely to be in higher categories of threat across Europe, particularly in the Mediterranean Sea where threat levels are generally higher (Figure 2a,b). When considering maximum body size in the Northeast Atlantic only, for every one unit increase in maximum body size (i.e. cm total length), the odds of a species being in an IUCN category of equal or higher threat increase by 0.98 (Figure 3a, Table S1). Similarly, in the Mediterranean Sea, for every one unit increase in maximum body size, the odds of a species being in an IUCN category of equal or higher threat increase by 0.94 (Figure 3a, Table S1). All other things being equal, a shark or ray of three metres total length in the Northeast Atlantic has a 71.7% probability of being in a threatened category (e.g. the Sandbar Shark, *Carcharhinus plumbeus*, Nardo 1827) compared to a 1.5 m species, which has a 39.4% probability of the same (e.g. the Angular Roughshark, *Oxynotus centrina*, Linnaeus 1758; Figure 2a). Whereas, in the Mediterranean Sea the Sandbar Shark is 84.5% likely to be in a threatened category and the Angular Roughshark is 62.1% likely to be threatened in this sub-region (Figure 2b). Hence, the conservation status of a 1.5 m shark or ray in the Mediterranean Sea is closer to that of a three metre species in the Northeast Atlantic, showing much less difference in likely conservation status between similar sized species in the Mediterranean Sea.

**Figure 2.**
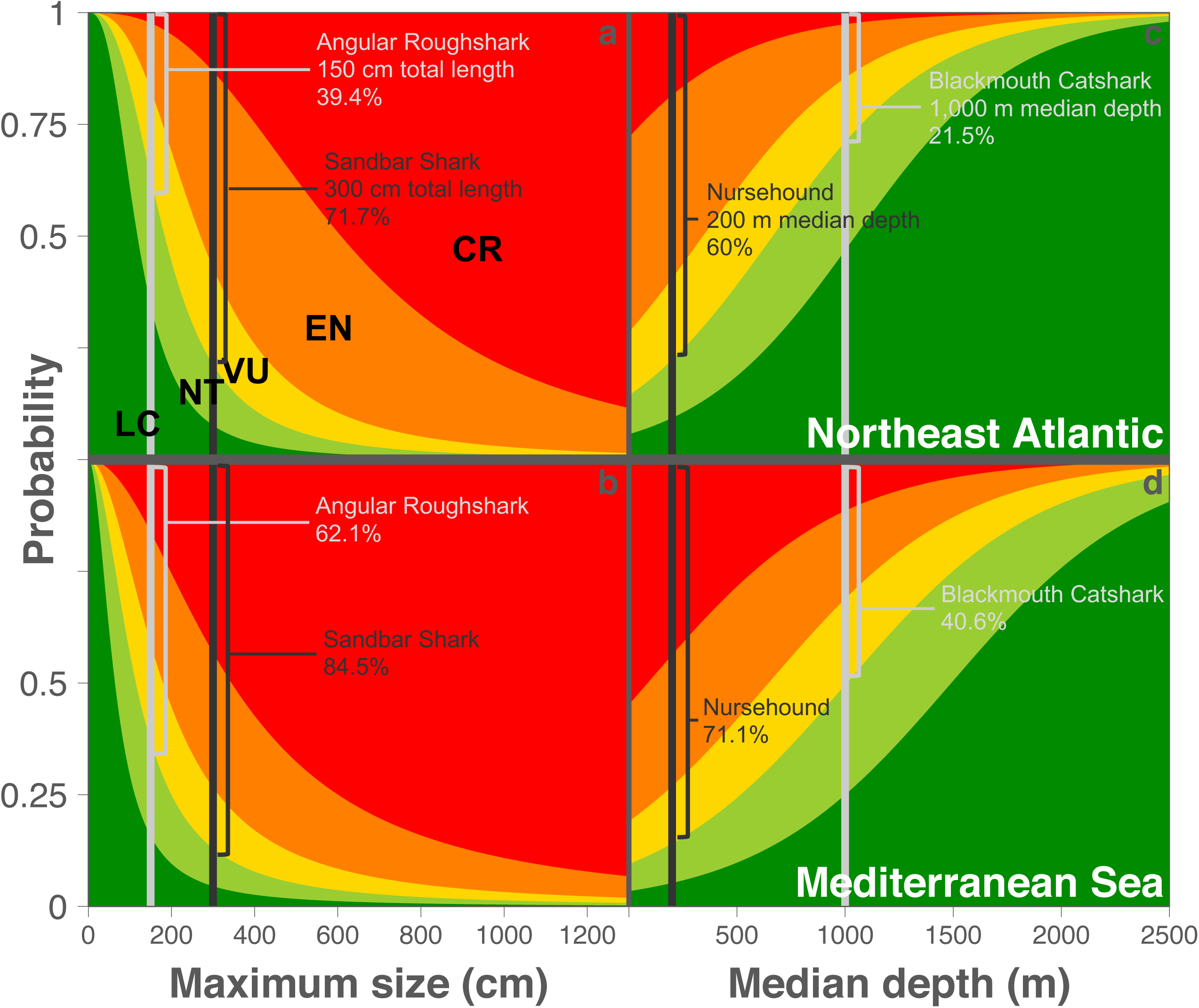
The effects of size and depth on shark and ray conservation status in Europe. Histograms of the probability of an evaluated shark or ray being listed as either Critically Endangered (CR), Endangered (EN), Vulnerable (VU), Near Threatened (NT), or Least Concern (LC) based on single-trait Cumulative Link Mixed-effects Model outputs for maximum body size (cm; panels a and b) and median depth (m; panels c and d). Data include all evaluated species (*n*= 94 Northeast Atlantic, panels a and c; and *n*=58 Mediterranean Sea, panels b and d) and exclude all Data Deficient species. Dark grey vertical bars indicate large (300 cm total length, a,b) or shallow (200 m median depth, c,d) species; light grey bars represent small (150 cm total length, a,b) or deep (1,000 m median depth, c,d) species. Brackets beside bars indicate the probability of each species being categorised as threatened (CR, EN, or VU) on the IUCN Red List.

**Figure 3.**
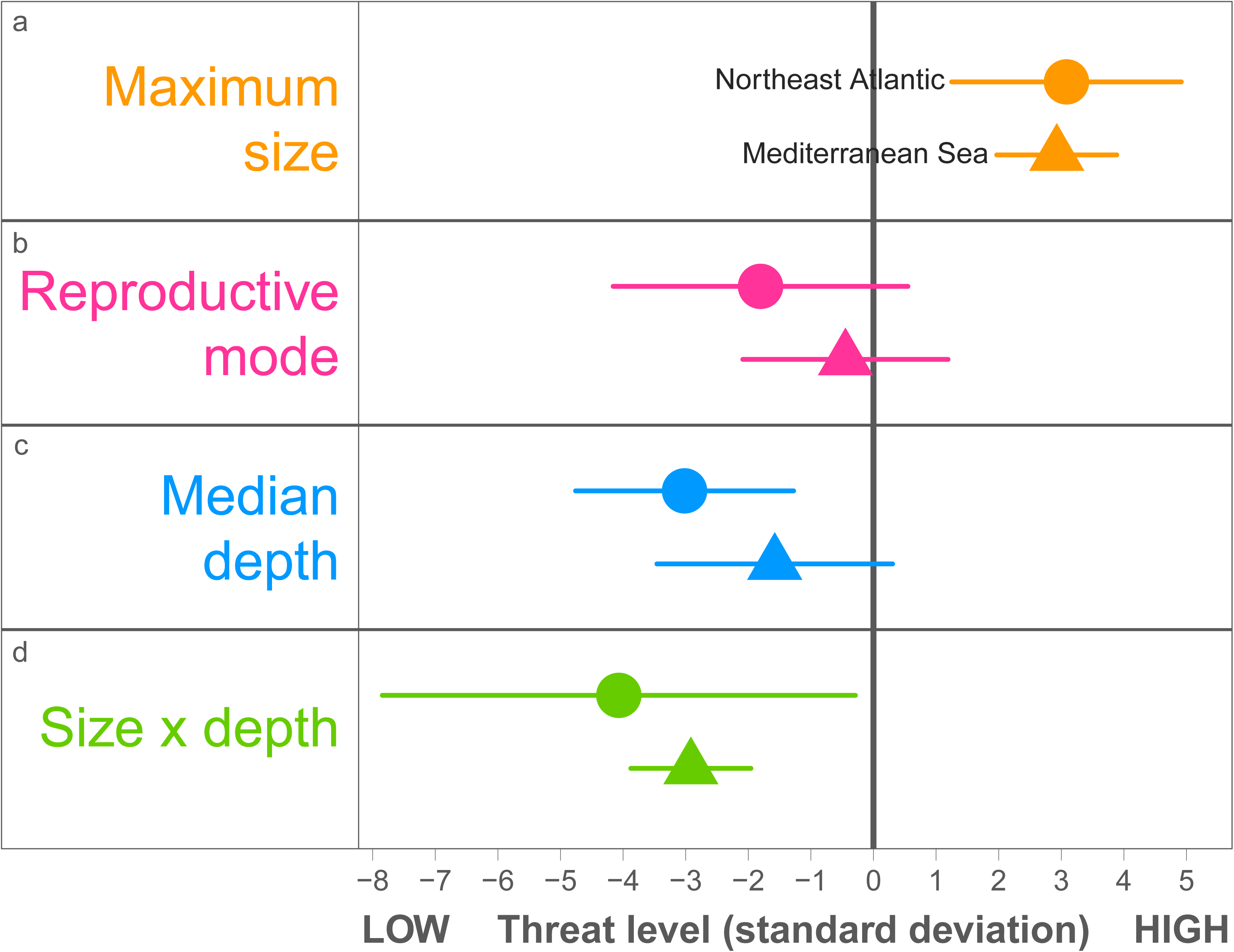
Effects of biological and ecological traits on Europe’s shark and ray conservation status. Standardized effect sizes with 95% confidence intervals. Cumulative link mixed effect models with maximum body size (a), reproductive mode (b), median depth (c), and the interaction between size and depth (d) as fixed effects and taxonomic Family as a random effect to account for phylogenetic non-independence. Circular and triangular points represent the best explanatory and predictive model for the Northeast Atlantic and Mediterranean Sea, respectively, which in both cases included maximum body size, reproductive mode, and the interaction between maximum size and median depth. Data for maximum size and median depth were centred and standardised by two standard deviations, while reproductive mode is a binary trait where oviparous species = 1 and viviparous species = 0.

Sharks and rays with greater depth distributions are more likely to be in lower categories of threat in the Northeast Atlantic (Figure 2c), but this pattern is muted in the Mediterranean Sea because threat levels are generally high for species across all depth distributions (Figure 2d). When considering median depth in the Northeast Atlantic only, for every one unit increase in median depth (i.e. metres), the odds of a species being in an IUCN category of higher threat decrease by 0.04 (Figure 3c, Table S1). In the Mediterranean Sea, for every one unit increase in median depth, the odds of a species being in a higher category of threat decrease by 0.24 (Figure 3c, Table S1). A similar-sized shark or ray with a median depth of 200 m has a 60% chance of being threatened in the Northeast Atlantic (e.g. the Nursehound, *Scyliorhinus stellaris*, Linnaeus 1758), compared with a species with a median depth of 1,000 m (e.g. the Blackmouth Catshark, *Galeus melastomus*, Rafinesque 1810), which has a 21.5% chance of being threatened in the same sub-region (Figure 2c). In the Mediterranean Sea the difference in risk is muted because of the greater reach of fisheries there: the Nursehound is 71.1% likely to be threatened, while the Blackmouth Catshark is 40.6% likely to be threatened in this sub-region (Figure 2d). Again, there is less differentiation between shallow and deepwater conservation status for Mediterranean species than Northeast Atlantic, and a higher likelihood of being threatened overall.

When maximum size, median depth, and reproductive mode are all considered, the odds of an egg-laying (oviparous) species being in a higher threat category decrease by 0.14 in the Northeast Atlantic (Figure 3b, Table S1). This effect was not significant in the Mediterranean Sea, again because the trait sensitivity is overridden or muted by the higher degree of exposure to fishing (Figure 3b, Table S1).

The most at-risk shark and ray species across Europe are therefore larger-bodied species restricted to the most heavily fished 0–400 m depth zone. The interaction between size and depth is such that for every unit increase in both size (cm) and depth (m), the odds of a shark or ray being in a higher category of threat decrease by 0.02 in the Northeast Atlantic, and by 0.05 in the Mediterranean Sea (Figure 3d, Table S1).

### 3.3 Predicted versus evaluated conservation status

The model with highest predictive accuracy (AUC) for both European sub-regions includes body size, reproductive mode, and the interaction between size and depth (Table 2, Figure 3). For the Northeast Atlantic, more than eight times out of ten, the top model predicts the correct category for the Critically Endangered, Endangered, Least Concern, or grouped threatened categories (Table S2). The Vulnerable category is predicted correctly more than six times out of ten, and the Near Threatened less than four times out of ten. The top model for the Mediterranean Sea predicts the Least Concern category correctly more than eight times out of then, and the Critically Endangered and Near Threatened categories more than seven times out of ten. Both sub-regional top models are weaker at predicting mid-range categories, with the Endangered category predicted correctly less than five times and the Vulnerable category less than six times out of ten (Table S2).

**Table 2.**
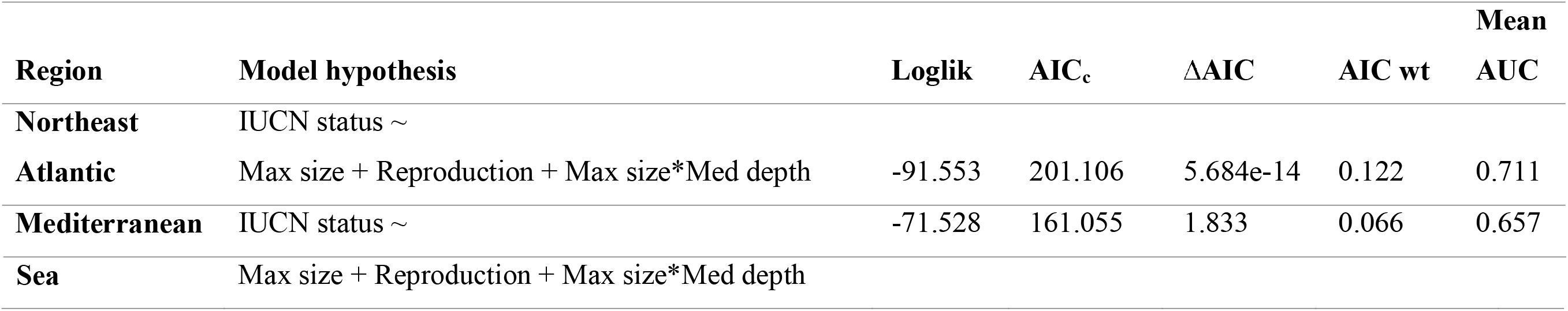
Summary of top Cumulative Link Mixed-effects Models for predicting IUCN status of Northeast Atlantic and Mediterranean sharks and rays. Models included all evaluated species (*n*=94 Northeast Atlantic and *n*=58 Mediterranean Sea). Top predictive models for both sub-regions with ΔAIC <2 included maximum size (cm), reproductive mode (oviparous=1, viviparous=0), and the interaction between maximum size and median depth (m). Maximum size and median depth were centred and standardised by two standard deviations. Each species was dropped one-at-a-time from the model and the IUCN status predicted. Comparison between evaluated and predicted statuses determined the predictive accuracy of each model. Model accuracy was measured as the Area Under the Curve (AUC) from the ROCR package in R version 3.5.2 (Sing et al., 2005) by scoring each category as one and all four other categories as zero to determine the predictive accuracy of all five separately (Critically Endangered, Endangered, Vulnerable, Near Threatened, Least Concern). To determine the top predictive model overall, the mean of all five category AUC values was calculated. Left to right: Loglik = log likelihood, AIC_c_ = AIC corrected for small sample size, ΔAIC = delta AIC, AIC wt = AIC weight, mean AUC = Area Under Curve averaged across the five AUC values for each IUCN category (see Table S2 for complete list of AUC values).

Almost half of Northeast Atlantic, and two-thirds of Mediterranean Data Deficient sharks and rays are predicted-to-be-threatened with an elevated risk of extinction (Figure 4c). This percentage of *predicted* threatened sharks and rays is greater than *evaluated* threat levels in the Northeast Atlantic (46% *predicted* versus 38% *evaluated* threatened), and similar to *evaluated* threatened in the Mediterranean Sea (67% *predicted* versus 66% *evaluated* threatened). The 12 Northeast Atlantic species predicted-to-be-threatened comprise 11 sharks and one ray (Figure 4a). All 12 Northeast Atlantic predicted-to-be-threatened species range from 89–640 cm total length, with depth ranges overlapping with fishing activity, and are viviparous. The ten predicted-to-be-threatened Mediterranean species comprise nine sharks and one ray (Figure 4b, Table S3). All nine species range from 114–427 cm total length, overlap significantly with the heavily fished depth zone, and are viviparous.

**Figure 4.**
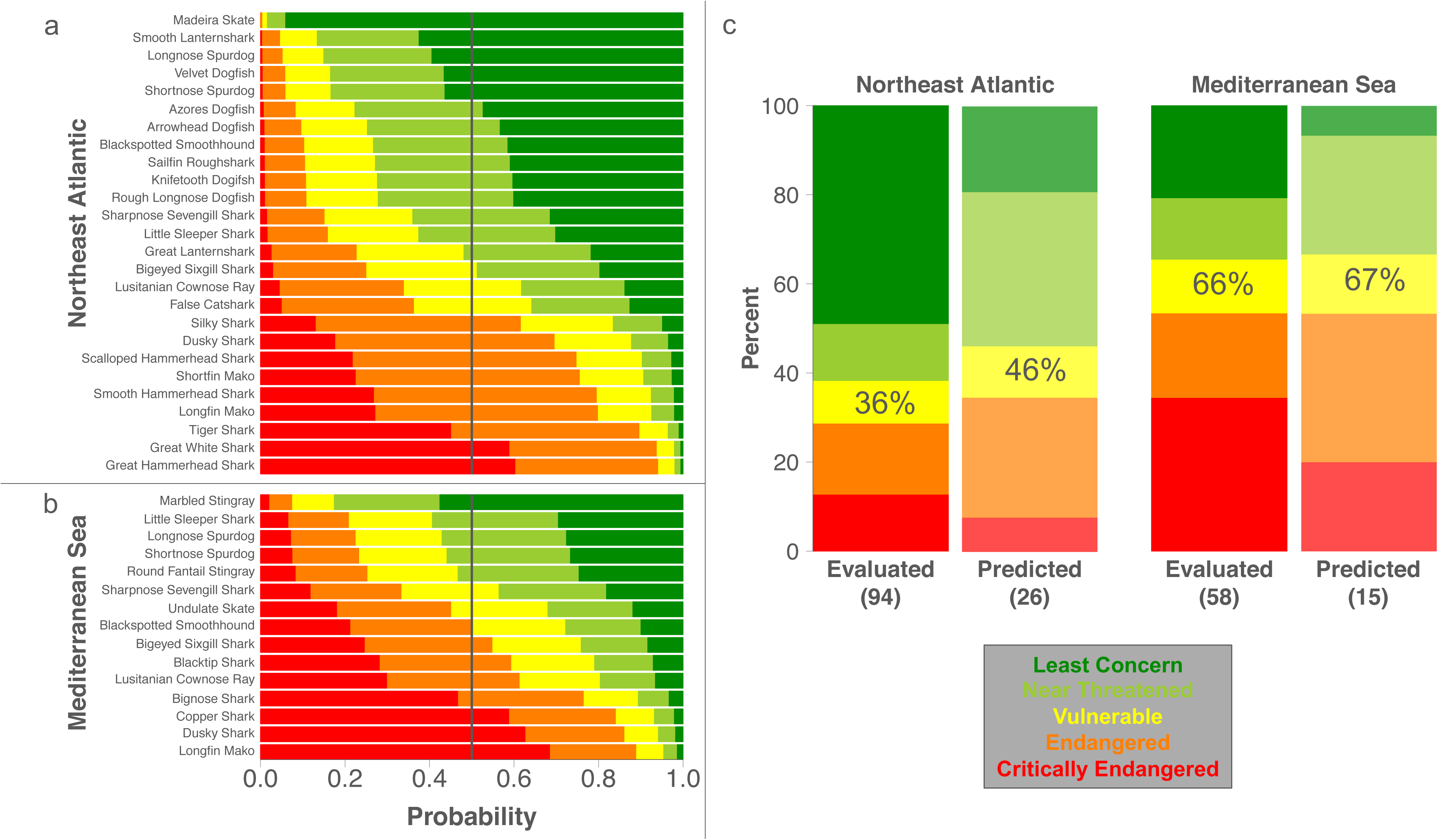
*Predicted* and *evaluated* conservation status of Europe’s sharks and rays. Top Cumulative Link Mixed-effectd Models including maximum body size, reproductive mode, and the interaction between size and median depth as fixed effects (for both sub-regions) and taxonomic Family as a random effect to account for phylogenetic non-independence. Panel a shows the probability of all 26 Data Deficient Northeast Atlantic species, and panel b of all 15 Mediterranean Sea species, being in each IUCN Red List category based on these top explanatory models. The vertical line cutting down panels a and b represents the 50% cut-off classification used to assign the final IUCN categorisations (according to the category bar the line crosses). Data for size and depth were centred and standardised by two standard deviations, while reproductive mode is a binary trait where oviparous species = 1 and viviparous species = 0. Panel c shows the proportion of sharks and rays in the Northeast Atlantic (left) and Mediterranean Sea (right) both *evaluated* and *predicted* to be in each IUCN category. Percentage values within each yellow bar indicate the total percentage of *evaluated* threatened and *predicted*-to-be-threatened species in each set, while numbers within brackets below each bar indicate the total number of species included in each set.

The distribution of both *evaluated* and *predicted* Northeast Atlantic listings in each case shows a median categorisation of Near Threatened, whereas in the Mediterranean Sea the median categorisation for both is Endangered (Figure S1). Overall, Northeast Atlantic and Mediterranean Sea listings have opposing distributions, with the majority of Northeast Atlantic species non-threatened and the majority of Mediterranean species threatened (Figure S1).

### 3.4 Europe’s most threatened Data Deficient sharks and rays

The species predicted to have the most elevated extinction risk (i.e. Critically Endangered) across Europe are all viviparous, large-bodied (349–640 cm total length) sharks whose median depths range from 40.5–350 m, hence overlapping greatly with the heavily fished zone (0–400 m depth; Table S3). In the Northeast Atlantic, the Great White Shark (*Carcharodon carcharias*, Linnaeus 1758) and Great Hammerhead Shark (*Sphyrna mokarran*, Rüppell 1837) are predicted to be Critically Endangered (Figure 4a; Table S3). While in the Mediterranean Sea, the Dusky Shark (*Carcharhinus obscurus*, Lesueur 1818), Copper Shark (*Carcharhinus brachyurus*, Günther 1870), and Longfin Mako (*Isurus paucus*, Guitart 1966) are predicted to be Critically Endangered (Figure 4b; Table S3). With this categorical regression approach, we identify a total of 14 Critically Endangered species in the Northeast Atlantic (approximately one third of threatened) and 23 Critically Endangered species in the Mediterranean Sea (approximately one half of threatened; Figure 5). If conservation efforts were focused on all imperilled species, i.e. the combined *evaluated*-threatened and *predicted*-to-be-threatened species, there would be a target list of 48 species to protect in the Northeast Atlantic and 48 species in the Mediterranean Sea (Table 3, Figure 5).

**Figure 5.**
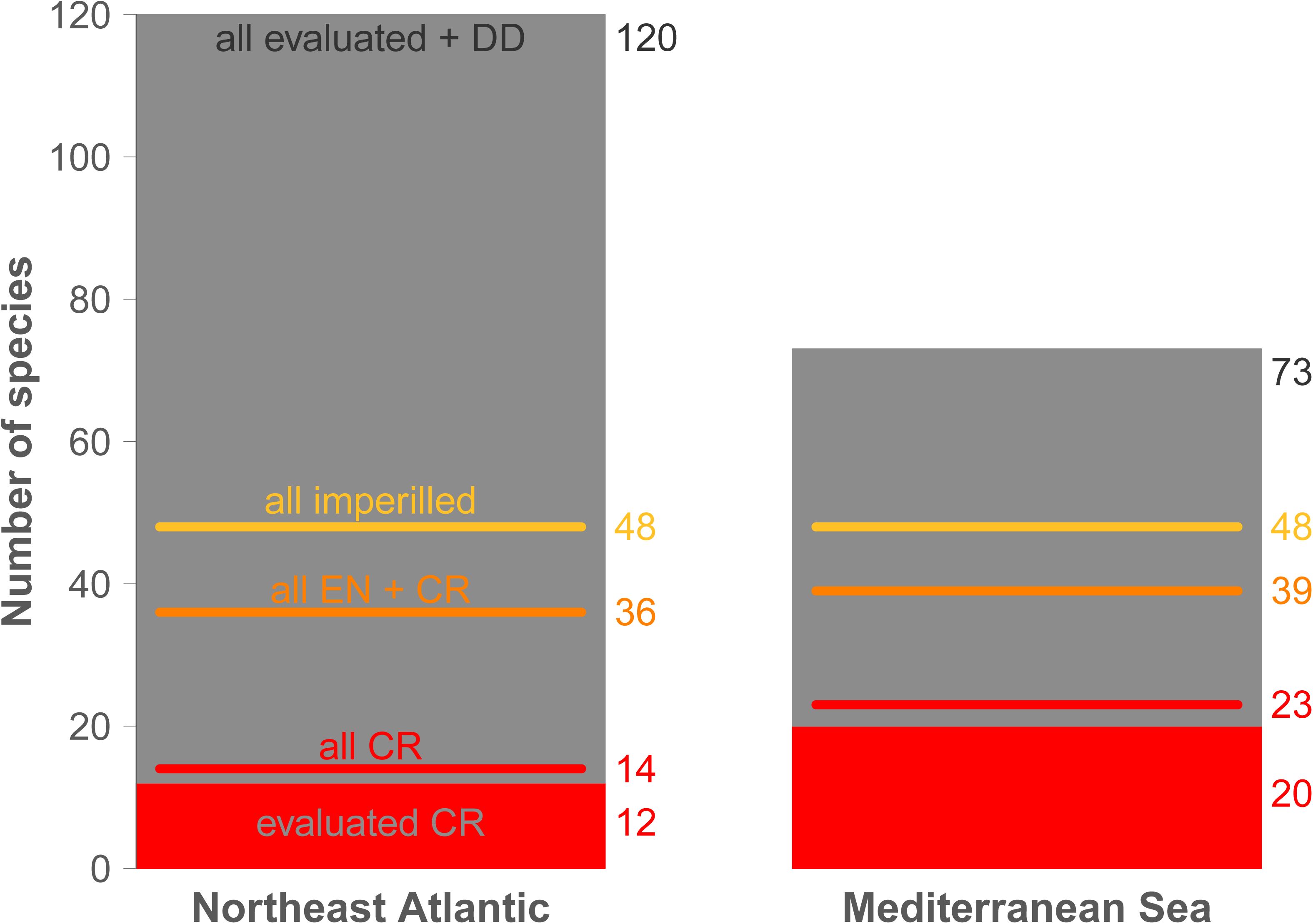
Informing shark and ray conservation efforts in Europe with categorical predictions. Solid grey bars represent species of all IUCN categories excluding those officially evaluated by the IUCN as Critically Endangered (CR), which are represented by solid red blocks. There are 120 shark and ray species in the Northeast Atlantic (left) and 73 in the Mediterranean Sea (right). Horizontal red lines indicate the addition of all Data Deficient species *predicted* to be Critically Endangered, to the *evaluated* block. Orange lines indicate all *evaluated* and *predicted*-to-be-Endangered and Critically Endangered species, while yellow lines show all imperilled (i.e. *evaluated* and *predicted*-to-be-threatened) species (Vulnerable, Endangered, and Critically Endangered). Numbers beside bars indicate total number of species within each relevant grouping.

**Table 3.**
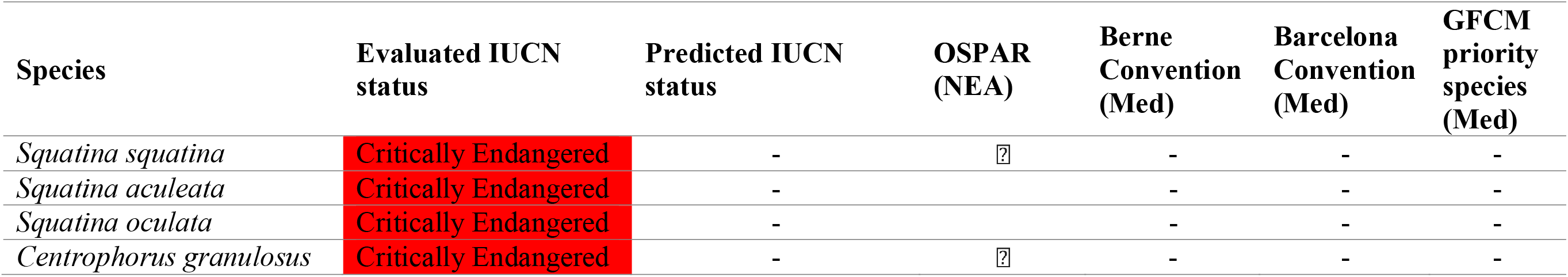
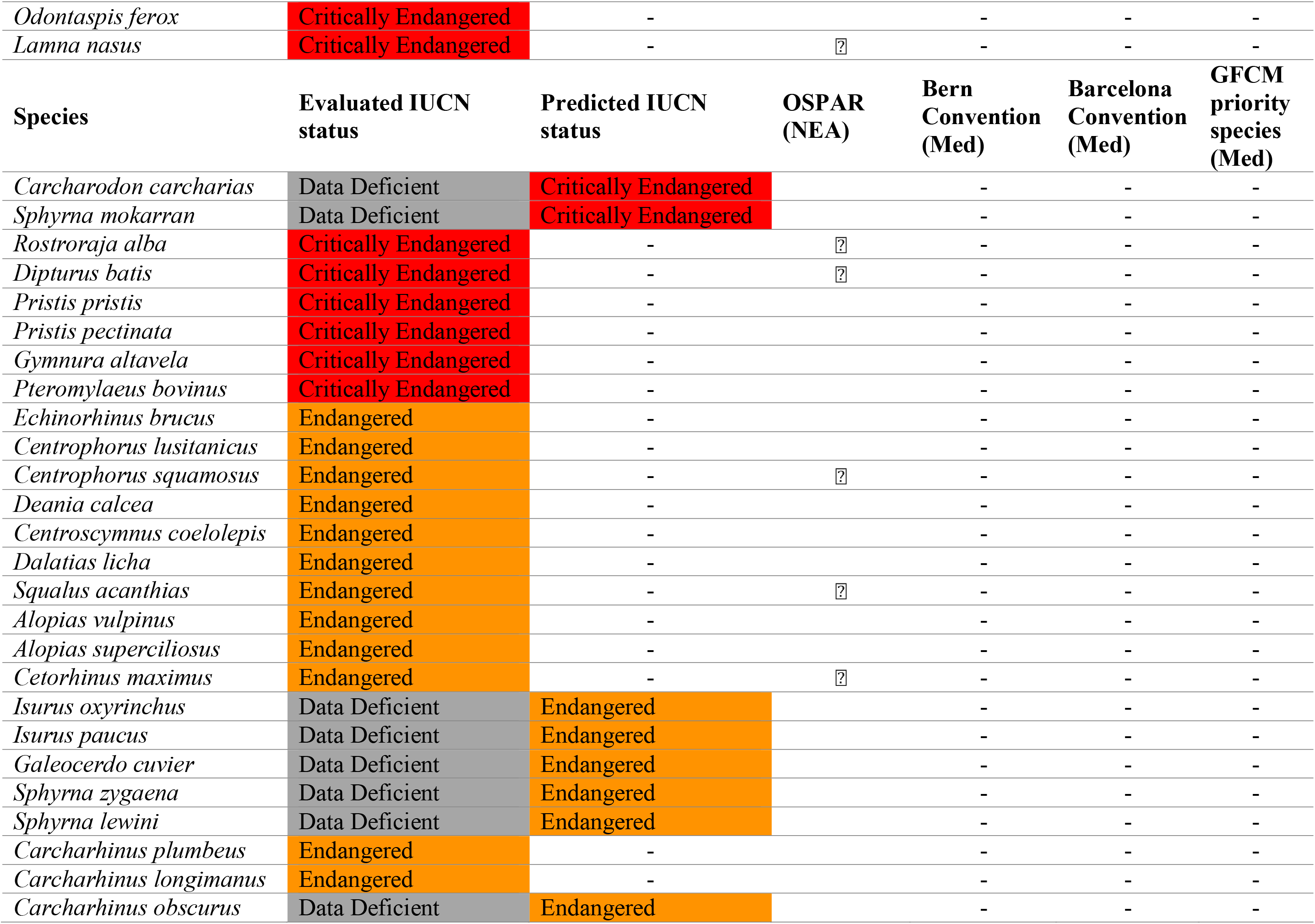
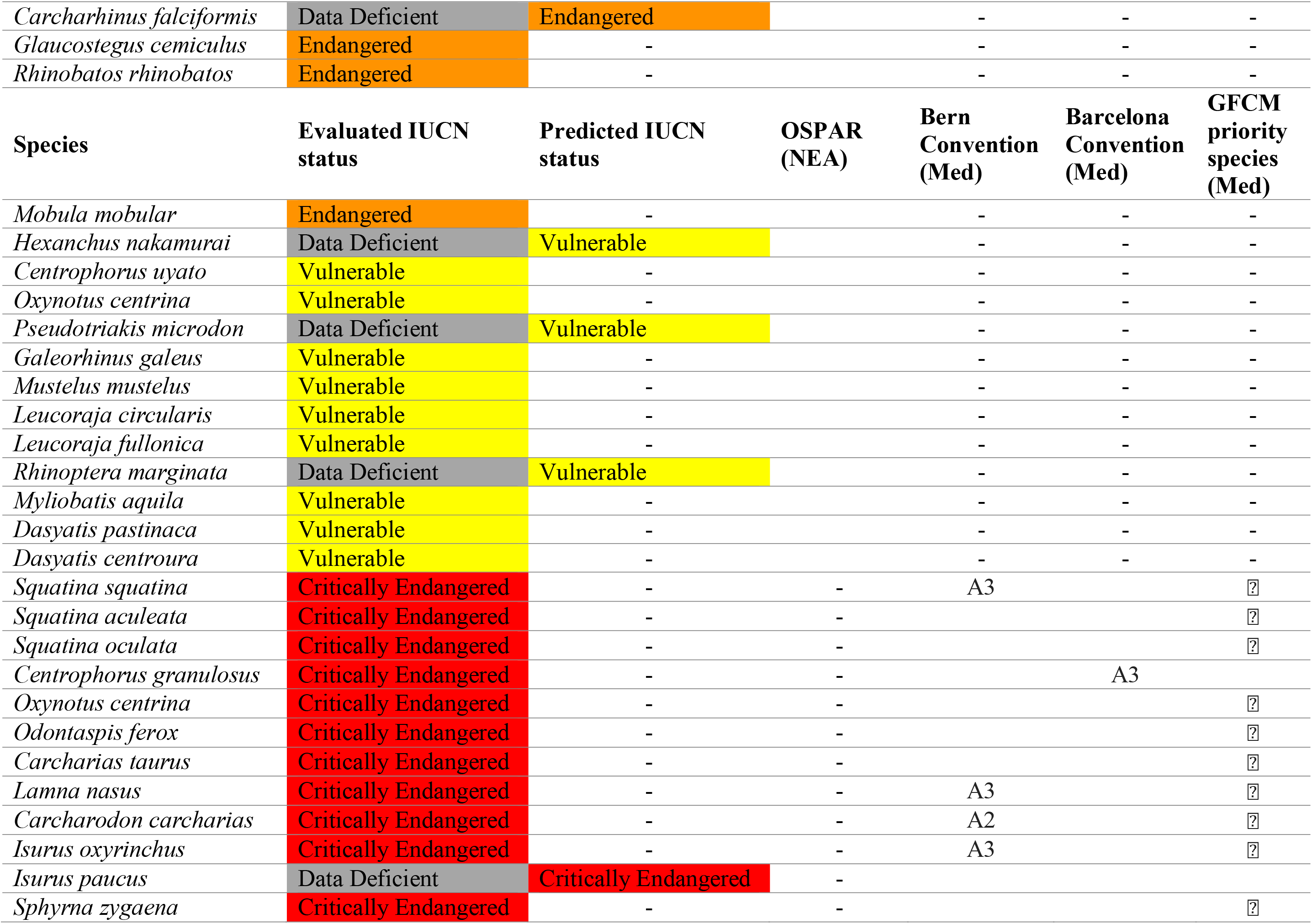
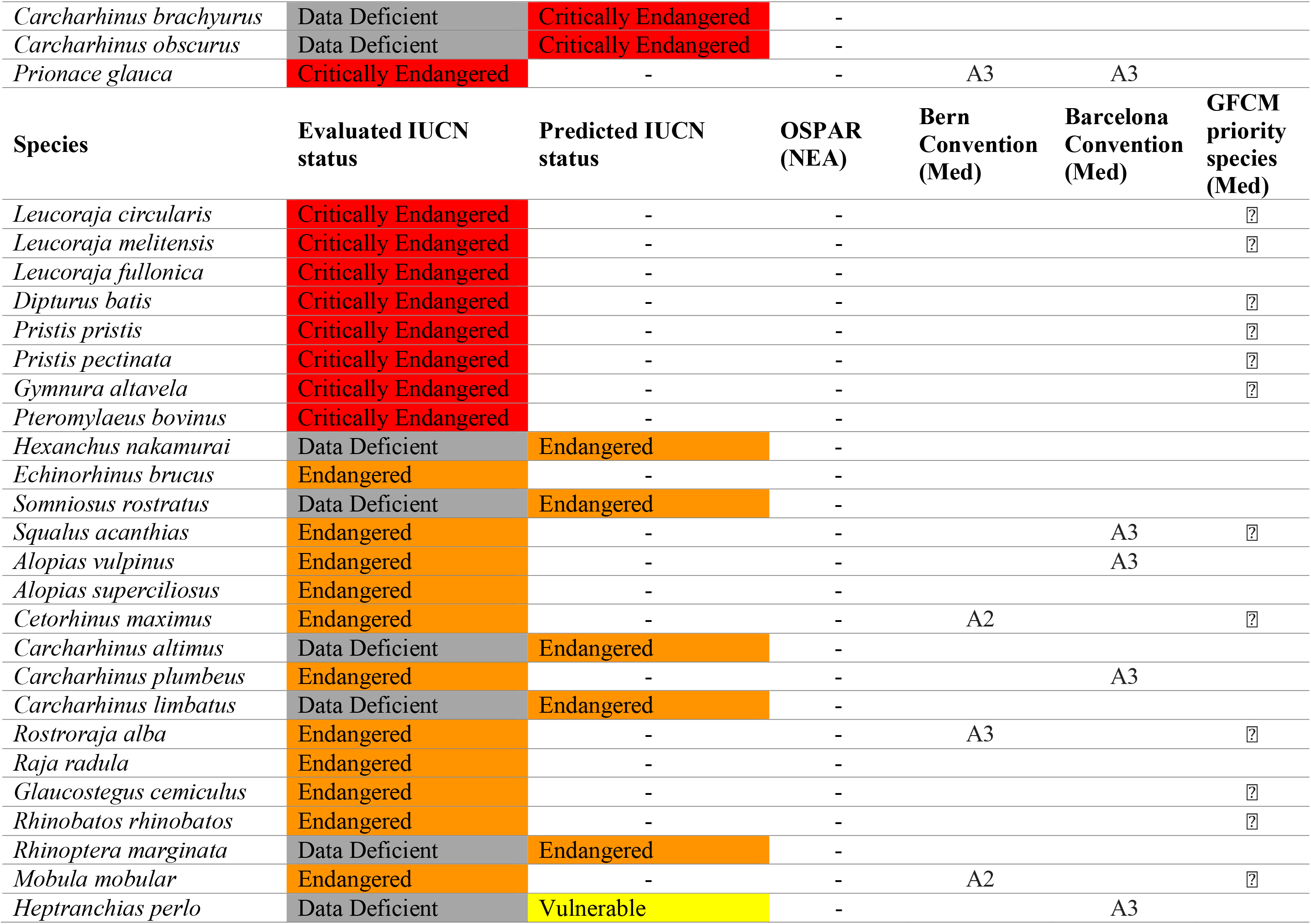
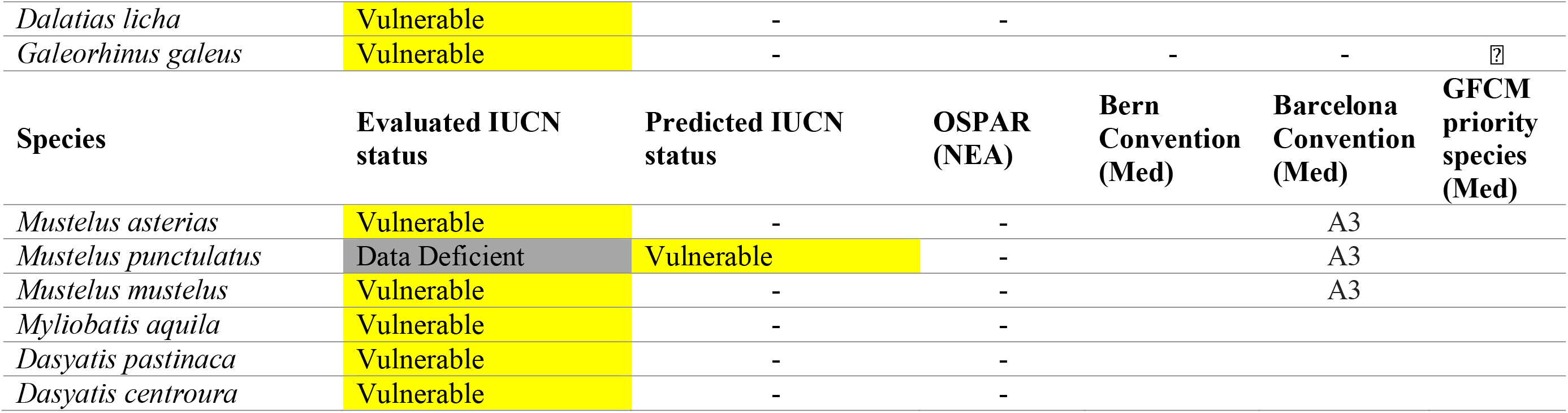
Current consideration of all imperilled sharks and rays in European waters by regional conventions and priority species lists. Relevant listings of all imperilled (i.e. *evaluated*-threatened and *predicted*-to-be-threatened) sharks and rays in Europe on regional and global protection-focused conventions, by major sub-region (Northeast Atlantic then Mediterranean Sea). Blanks indicate no listing, while hyphens indicate inapplicability of a convention to a species within a certain sub-region. Where a convention has multiple appendices, the applicable appendix number is indicated (e.g. A2, A3) instead of a tick mark. Species are listed taxonomically within each threatened IUCN category – Critically Endangered (CR), Endangered (EN), and Vulnerable (VU) – in descending order of threat. Conventions left to right: Oslo-Paris Convention (OSPAR; applicable to Northeast Atlantic Ocean only); Berne Convention (applicable to Mediterranean Sea only); Barcelona Convention (Mediterranean Sea only); and the General Fisheries Commission for the Mediterranean (GFCM) priority species list (Mediterranean Sea only). The Great Hammerhead Shark (*Sphyrna mokarran*) is currently one of 24 species included on the GFCM priority species list, but has not been included in the Mediterranean section of this table as it is now considered a Vagrant species in the Mediterranean Sea, as per IUCN definition (IUCN, 2012a).

## 4 DISCUSSION

### 4.1 Regional versus global IUCN Red List Status

Here, we show that sharks and rays are proportionally more threatened in the two main sub-regions of Europe than the global reported threat rate, particularly when we account for the predicted risk status of Data Deficient species. Overall, we estimate that there are 40% (48) imperilled species (*evaluated* threatened and *predicted*-to-be-threatened) in the Northeast Atlantic and 67% (48) in the Mediterranean Sea. Compared with other vertebrate groups, these threat levels not only exceed those of global sharks and rays (23.9%, *n*=249 of 1,041, (Dulvy et al., 2014), but also that of amphibians: the most imperilled assessed group to date (41%, *n*=2,561 of 6,284, Hoffmann et al., 2010). Furthermore, whereas almost half of global sharks and rays are Data Deficient (46.8%, *n*=487), approximately one-fifth of Europe’s species are Data Deficient, which is also closer to global amphibian data deficiency proportionally (26%, *n*=1,597; Hoffmann et al., 2010). The high levels of threat and relatively low levels of data deficiency in Europe result from the region’s comparably long-standing history of fishing and data collection compared with the rest of the world (Barrett et al., 2004; Hoffmann, 1996). We next consider: (1) the biological and ecological traits driving these regional threat levels, (2) the differences between *evaluated* and *predicted* conservation status, and (3) how categorically predicting such could help narrow the focus of conservation efforts overall.

### 4.2 Biological and ecological predictors of conservation status

Sharks and rays with both larger maximum body size and shallower depth distribution are more likely to face an elevated risk of extinction than smaller (faster-growing) species that live predominantly in deeper water. Fishing is the greatest threat to sharks and rays (McClenachan, et al., 2012), and it is greatest from 0–400 m depth, but in European waters lower levels of fishing activity occur down to at least 1,000 m (Amoroso et al., 2018; Morato et al., 2006). In waters deeper than the reach of fisheries, a species can be very large-bodied and not threatened at all, because body size has little influence over conservation status unless it is combined with a major threat (Fernandes et al., 2017; Owens & Bennett, 2000; Reynolds et al., 2005a). For example, the Goblin Shark (*Mitsukurina owstoni*, Jordan 1898) reaches 617 cm total length with a depth range of 40–1,569 m. This deepwater shark is listed as Least Concern in the Northeast Atlantic as the majority of its depth range offers refuge from fishing activity. By contrast, the Common Thresher Shark (*Alopias vulpinus*, Bonnaterre 1788) reaches 573 cm total length, has a depth range of 0–366 m (i.e. entirely overlapping with the heavily fished zone), and is listed as Endangered in the same sub-region. Large body size has been associated with increased probability of extinction for numerous taxonomic groups (e.g. Cardillo et al., 2011; Comeros-Raynal et al., 2016; Field et al., 2009). Large body size is known to be correlated to a slow speed-of-life, but also as an impediment to evading capture in fishing gear. For shark and ray conservation, accounting for this relationship between size and susceptibility to capture is complicated by the issue of bycatch. More sharks and rays are threatened by incidental catch than by actual target fisheries (Dulvy et al., 2014), where their higher intrinsic sensitivity is not accounted for by fisheries management regimes that are focused on faster growing, less sensitive teleost (bony fish) species. Across Europe, the predominant fishing techniques are highly unselective, such as multi-species trawling (Smith & Garcia, 2014). The incentive for fishers to increase selectivity to benefit non-target species is low when this action would undoubtedly coincide with reduced target catch. The consequent unselective fishing of non-target species is a major driver of the high threat levels among Europe’s sharks and rays and could lead to overlooked local extinctions. This predicament alone presents incentive to better understand the status of Data Deficient species, particularly in heavily fished waters such as Europe.

The conservation status of sharks and rays in the Mediterranean Sea appears much worse than the Northeast Atlantic, which can partly be explained by the lack of depth refuge for sharks and rays from heavy fishing activity in this sub-region. The Mediterranean Sea has a longer history of fishing than the Northeast Atlantic, and nowadays, that fishing is not managed as efficiently as it is in the Northeast Atlantic (Fernandes et al., 2017; Smith & Garcia, 2014). A semi-enclosed sea equates to many more sites for landing catches, none of which are being consistently monitored. Further exacerbating this lack of monitoring, the Mediterranean fishery principally comprises higher numbers of smaller artisanal vessels, compared with fewer, more readily trackable commercial vessels in the Northeast Atlantic (Smith & Garcia, 2014). Semi-enclosed seas are also more susceptible to other major threatening events than open oceans, such as ocean acidification, rising temperatures, and coastal pollution and development (Caddy, 2000). Despite these logical contributors to the higher threat levels seen among Mediterranean sharks and rays, the difference in conservation status between both major European sub-regions can largely be attributed to the differing taxonomic and hence trait composition (120 Northeast Atlantic and 73 Mediterranean Sea sharks and rays). There are 35 deepwater shark and ray species that exist in the Northeast Atlantic exclusively outside the reach of fisheries, and are all therefore listed as Least Concern, which do not occur in the Mediterranean Sea. If those species are removed from the Northeast Atlantic species list, we see the same number of imperilled species in each sub-region, and a much more similar overall proportion of threat (Table S4). Meanwhile, the median depths of all 73 Mediterranean Sea species overlap to some degree (if not entirely) with the heavily fished 0–400 m depth zone. This explains why median depth, and the interaction between maximum body size and median depth, are weaker explanatory variables in this sub-region: depth refuge from fishing activity simply does not exist for sharks and rays in the Mediterranean Sea.

Perhaps the prevailing threat resultant from lacking refuge in the Mediterranean Sea also explains to some extent why oviparous species are not significantly lower-risk in this sub-region than viviparous species. Oviparity is characteristically associated with faster population growth rates than viviparity (Field et al., 2009). All of the most threatened sharks and rays in Europe are viviparous, likely because they are less able to withstand fishing pressure as effectively as typically faster-growing, egg-laying species.

### 4.3 *Predicted* versus *evaluated* conservation status

To provide taxon-wide estimates of extinction risk in the face of uncertainty, the IUCN assumes that the fraction of threatened Data Deficient species is the same as the proportion of *evaluated*- threatened species. While pragmatic, this is an assumption to be tested. A recent estimate of which Data Deficient sharks and rays might be classified as Least Concern or threatened revealed 14% of Data Deficient species were *predicted*-to-be-threatened (*n*=68 of 487), and overall 17.8% were *evaluated*-threatened (Dulvy et al., 2014). Taken together, there is an overall estimated global threat level of 23.9% imperilled sharks and rays (Dulvy et al., 2014). By comparison, this is much lower than the IUCN equal ratio approach, which hence yields an inflated estimate of 33% of sharks and rays threatened (Hoffmann et al., 2010).

This 1:1 ratio of *predicted*-to-*evaluated* threatened species proportions holds true for global birds (Class Aves), in which knowledge is significantly greater than for other taxa and hence there are few Data Deficient species (0.6%, *n*=63 of 10 500; Butchart & Bird, 2010). This 1:1 ratio approach, however, yields a 50% underestimation of globally Data Deficient threatened mammals, where one-third of *evaluated* species are threatened, whereas two-thirds of Data Deficient species are *predicted*-to-be-threatened (1:2, Jetz & Freckleton, 2015). In the case of mammals, reliance on this ratio could have devastating implications for species extinction rates by overlooking 50% of the recently predicted-to-be-threatened Data Deficient species. Conversely, using the 1:1 ratio of *evaluated* to *predicted* threatened status would overestimate globally Data Deficient threatened groupers, for which the proportion of *evaluated* threat is actually three times higher than that of *predicted* threat (3:1, Luiz et al., 2016).

Northeast Atlantic sharks and rays have a similar threat distribution, and hence, negative conservation implications to global mammals if the IUCN’s 1:1 ratio were to be relied upon (1:1.2 *evaluated* to *predicted* threat). Global sharks and rays have the opposite pattern, whereby conservation resources might be wasted on the protection of Data Deficient species according to this ratio. Yet, despite the high levels of data deficiency among Mediterranean sharks and rays compared with global birds, the present study shows this 1:1 ratio to be appropriate for this sub-regional taxonomic group also. The inconsistency in these risk ratio patterns across taxonomic groups, and geographic regions within taxonomic groups, highlights the need for taxon-specific predictions of threat among Data Deficient listings.

### 4.4 Updating protected species lists in Europe

There are a number of lists that flag species for protection, but many of these are now out-of-date. Ideally, all of the imperilled sharks and rays in Europe would be listed in the appendices of the appropriate conservation-focused conventions in the region, and their exploitation monitored and managed accordingly. In reality, only eight of the 48 imperilled species identified here are listed on the Oslo-Paris convention in the Northeast Atlantic (Table 3). While of the 48 imperilled species identified here in the Mediterranean Sea, only three (and five) are listed on Appendix II (and Appendix III) of the Berne Convention, nine on Appendix III of the Barcelona Convention, and 23 on the General Fisheries Commission for the Mediterranean priority species list (Table 3). The Great Hammerhead Shark is currently one of the 24 species included on the GFCM priority species list for the Mediterranean Sea, but since the 2015 European Red List reassessment this species is considered a Vagrant in this basin. We therefore consider only 23 species on the GFCM priority species list in this study (Table 3). Clearly, there is significant scope to update these lists to ensure protection of all imperilled species in both sub-regions.

### 4.5 Incorporating predictions into a Red List Index for 2020 target tracking

Categorical predictions of IUCN status enable the inclusion of Data Deficient species in aggregate species conservation status analyses, from which they are currently excluded for all taxonomic groups. The Convention on Biological Diversity’s Aichi Targets are monitored using indicators, such as the IUCN’s Red List Index, which is an indicator of the change in aggregate extinction risk over time (Brooks et al., 2015). Predicting IUCN status for Data Deficient species enables their addition to such indices, which would in turn give conservation planners a more holistic idea of conservation status. With sufficient model accuracy, it is likely more informative to include these predictions in such indices than to exclude them altogether. Upon completion of the ongoing global reassessment of sharks and rays, this methodology can be extrapolated to the global dataset for inclusion in the global Red List Index. This approach would prove even more accurate for highly data-sufficient groups such as birds. Resource limitations have hindered scientists and conservationists from focusing on Data Deficient species historically, but categorical predictions of conservation status are a cost-effective solution to this shortcoming (Bland et al., 2015), at least until data availability and resources allow for fully comprehensive IUCN assessment of these species. This case study, and the extrapolation to the highly Data Deficient global shark and ray dataset, will ideally be the first step towards applying this predictive approach to some more Data Deficient groups, such as plants and invertebrates.

## Supporting information

SOM

## ACKNOWLEDGEMENTS

We thank the IUCN Species Survival Commission Shark Specialist Group members, staff, and volunteers, and the IUCN head office for their logistical support throughout the Red List assessment process. In particular, we thank Alvaro Abella, David Allen, Michel Bariche, Tom Blasdale, Mohamed N Bradaï, Elena Buscher, Maurice Clarke, Simona Clo, Andrey Dolgov, Manuel Dureuil, Jim Ellis, Edward Farrell, Francesco Ferretti, Sonia Fordham, Sarah Fowler, Karen Frazer, Mariana García, Javier Guallart, Lucy Harrison, Ali Hood, Samuel Iglésias, Armelle Jung, James Kemp, Pete Kyne, Julia Lawson, Sophy McCully, Ana Nieto, Giuseppe Notarbartolo di Sciara, Caroline Pollock, Gina Ralph, Fabrizio Serena, Bernard Seret, Alen Soldo, Matthias Stehmann, Heike Zidowitz. We thank Tobias Sing, Dan Greenberg, Nathan Pacoureau, Chris Brown, and Rylee Murray for statistical advice, and members of the Dulvy Lab, Earth_2_Ocean Lab and Stats Beerz for comment on drafts and statistical advice.

