## Supplementary material for "Predicting the conservation status of Europe’s Data Deficient sharks and rays": SOM

**Supplementary Table A1**

Summary of predictive cumulative link mixed effect models for biological and ecological correlates of IUCN classification for Europe’s two major sub-regions, the Northeast Atlantic and Mediterranean Sea. Data Deficient species were excluded from these ‘training’ models, which were based on all European sharks and rays evaluated by the IUCN to use subsequently for predicting the IUCN statuses of the Data Deficient species. Species were categorised under five IUCN categories: Critically Endangered, Endangered, Vulnerable, Near Threatened, and Least Concern, for *n*=94 Northeast Atlantic and *n*=58 Mediterranean Sea species.

|  | Parameter | Coefficient | Standard Error | z value | p value | Degrees of freedom (*k*) |
| --- | --- | --- | --- | --- | --- | --- |
| *Northeast Atlantic*  Log likelihood:  -91.55305  AIC: 201.1061 | Maximum size | 3.0832 | 0.9356 | 3.2954 | 0.001 | 93 |
|  | Reproductive mode | –1.8040 | 1.2011 | –1.5020 | 0.133 | 93 |
|  | Median depth | –3.0177 | 0.8883 | ­–3.3972 | 0.001 | 93 |
|  | Max. size * med. depth | –4.0675 | 1.9276 | –2.1102 | 0.035 | 93 |
| *Mediterranean Sea*  Log likelihood:  -71.52752  AIC: 161.055 | Maximum size | 2.9279 | 0.8362 | 3.5012 | 0.001 | 57 |
|  | Reproductive mode | –0.4480 | 0.9603 | –0.4666 | 0.641 | 57 |
|  | Median depth | –1.5772 | 0.6377 | –2.4734 | 0.013 | 57 |
|  | Max. size * med. depth | –2.9165 | 1.4647 | –1.9911 | 0.046 | 57 |

**Supplementary Table A2**

Predictive model accuracy for each top model from both major European sub-regions. Both top models included the IUCN category as the response variable, maximum body size, reproductive mode, and the interaction between maximum size and median depth as fixed effects, and taxonomic Family as a random effect to account for phylogenetic non-independence. Predictive accuracy was determined using the Receiver Operating Characteristic curve R package, ROCR (Sing et al., 2005), which operates for binary classification only. Hence, each category was individually scored as one against the four remaining categories scored as zero to discern the accuracy for each category separately. The Area Under the Curve (AUC) score represents the probability of a model predicting the correct category, therefore scores closer to 1 represent high accuracy and those closer to zero indicate low predictive accuracy. CR = Critically Endangered. To choose the model with the highest overall predictive accuracy across categories, the mean AUC of all five categories was calculated.

|  | AUC | | |
| --- | --- | --- | --- |
| IUCN Category/grouping | | Northeast Atlantic: Model 1 | Mediterranean Sea: Model 4 |
| Critically Endangered | | 0.832 | 0.712 |
| Endangered | | 0.817 | 0.474 |
| Vulnerable | | 0.686 | 0.580 |
| Near Threatened | | 0.395 | 0.703 |
| Least Concern | | 0.825 | 0.815 |
| threatened | | 0.893 | 0.862 |
| Mean (excl. threatened) | | 0.711 | 0.657 |

**Supplementary Table A3**

Predicted IUCN categories of all Data Deficient sharks and rays in Europe by major sub-region. There are 26 Data Deficient sharks and rays in the Northeast Atlantic sub-region and 15 in the Mediterranean Sea. Predictions were made with cumulative link mixed effects models of biological and ecological traits of each species based on the known IUCN categorisations of all evaluated sharks and rays in Europe. Taxonomic Family was included in each model as a random effect to account for phylogenetic covariation. Final categorisations were allocated using the default probability cut-off value of 50%.

| Species Latin Name | Species Common Name | Predicted IUCN status | Maximum size (cm) | Median depth (m) | Reproductive mode |
| --- | --- | --- | --- | --- | --- |
| Northeast Atlantic Ocean | | |  |  |  |
| *Carcharodon carcharias* | Great White Shark | Critically Endangered | 640 | 126 | Viviparous |
| *Sphyrna mokarran* | Great Hammerhead Shark | Critically Endangered | 610 | 40.5 | Viviparous |
| *Isurus oxyrinchus* | Shortfin Mako | Endangered | 400 | 251 | Viviparous |
| *Isurus paucus* | Longfin Mako | Endangered | 427 | 180 | Viviparous |
| *Galeocerdo cuvier* | Tiger Shark | Endangered | 550 | 175.5 | Viviparous |
| *Sphyrna zygaena* | Smooth Hammerhead Shark | Endangered | 400 | 101 | Viviparous |
| *Sphyrna lewini* | Scalloped Hammerhead Shark | Endangered | 370 | 138 | Viviparous |
| *Carcharhinus obscurus* | Dusky Shark | Endangered | 349 | 200.5 | Viviparous |
| *Carcharhinus falciformis* | Silky Shark | Endangered | 316 | 259 | Viviparous |
| *Hexanchus nakamurai* | Bigeyed Sixgill Shark | Vulnerable | 180 | 355.5 | Viviparous |
| *Pseudotriakis microdon* | False Catshark | Vulnerable | 296 | 995 | Viviparous |
| *Rhinoptera marginata* | Lusitanian Cownose Ray | Vulnerable | 200 | 50.5 | Viviparous |
| *Heptranchias perlo* | Sharpnose Sevengill Shark | Near Threatened | 140 | 650 | Viviparous |
| *Deania hystricosa* | Rough Longnose Dogfish | Near Threatened | 111 | 885.5 | Viviparous |
| *Deania profundorum* | Arrowhead Dogfish | Near Threatened | 97 | 1030 | Viviparous |
| *Somniosus rostratus* | Little Sleeper Shark | Near Threatened | 143 | 1161 | Viviparous |
| *Scymnodalatias garricki* | Azores Dogfish | Near Threatened | 80 | 1150 | Viviparous |
| *Oxynotus paradoxus* | Sailfin Roughshark | Near Threatened | 118 | 492.5 | Viviparous |
| *Scymnodon ringens* | Knifetooth Dogfish | Near Threatened | 110 | 900 | Viviparous |
| *Etmopterus princeps* | Great Lanternshark | Near Threatened | 89 | 2425 | Viviparous |
| *Mustelus punctulatus* | Blackspotted Smoothhound | Near Threatened | 122 | 150.5 | Viviparous |
| *Zameus squamulosus* | Velvet Dogfish | Least Concern | 69 | 1000 | Viviparous |
| *Squalus Blainville* | Longnose Spurdog | Least Concern | 90 | 346.5 | Viviparous |
| *Squalus megalops* | Shortnose Spurdog | Least Concern | 94 | 366.5 | Viviparous |
| *Etmopterus pusillus* | Smooth Lanternshark | Least Concern | 50 | 1136 | Viviparous |
| *Raja maderensis* | Madeira Skate | Least Concern | 80 | 76 | Oviparous |
| Mediterranean Sea | | |  |  |  |
| *Isurus paucus* | Longfin Mako | Critically Endangered | 427 | 180 | Viviparous |
| *Carcharhinus brachyurus* | Copper Shark | Critically Endangered | 350 | 50.5 | Viviparous |
| *Carcharhinus obscurus* | Dusky Shark | Critically Endangered | 349 | 200.5 | Viviparous |
| *Hexanchus nakamurai* | Bigeyed Sixgill Shark | Endangered | 180 | 355.5 | Viviparous |
| *Somniosus rostratus* | Little Sleeper Shark | Endangered | 143 | 1161 | Viviparous |
| *Carcharhinus altimus* | Bignose Shark | Endangered | 282 | 221 | Viviparous |
| *Carcharhinus limbatus* | Blacktip Shark | Endangered | 193 | 15.5 | Viviparous |
| *Rhinoptera marginata* | Lusitanian Cownose Ray | Endangered | 200 | 50.5 | Viviparous |
| *Heptranchias perlo* | Sharpnose Sevengill Shark | Vulnerable | 140 | 650 | Viviparous |
| *Mustelus punctulatus* | Blackspotted Smoothhound | Vulnerable | 122 | 150.5 | Viviparous |
| *Squalus blainville* | Longnose Spurdog | Near Threatened | 90 | 346.5 | Viviparous |
| *Squalus megalops* | Shortnose Spurdog | Near Threatened | 94 | 366.5 | Viviparous |
| *Raja undulata* | Undulate Skate | Near Threatened | 114 | 101 | Oviparous |
| *Taeniurops grabata* | Round Fantail Stingray | Near Threatened | 100 | 50.5 | Viviparous |
| *Dasyatis marmorata* | Marbled Stingray | Least Concern | 60 | 38.5 | Viviparous |

**Supplementary Table A4**

Percent listings of sharks and rays under each IUCN category in each major European sub-region, indicating the similarity between the percent threatened species when Northeast Atlantic-exclusive deepwater species are removed from the species list and the remaining species list compared with the Mediterranean Sea.

| IUCN Category | Percent species listings | |
| --- | --- | --- |
|  | Northeast Atlantic | Mediterranean Sea |
| Critically Endangered | 10 | 27 |
| Endangered | 13 | 15 |
| Vulnerable | 8 | 10 |
| Near Threatened | 10 | 11 |
| Least Concern | 38 | 16 |
| Data Deficient | 22 | 21 |
| Percent threatened | 30 | 52 |
| Percent threatened (minus 35 deepwater NEA species) | **42** | **52** |

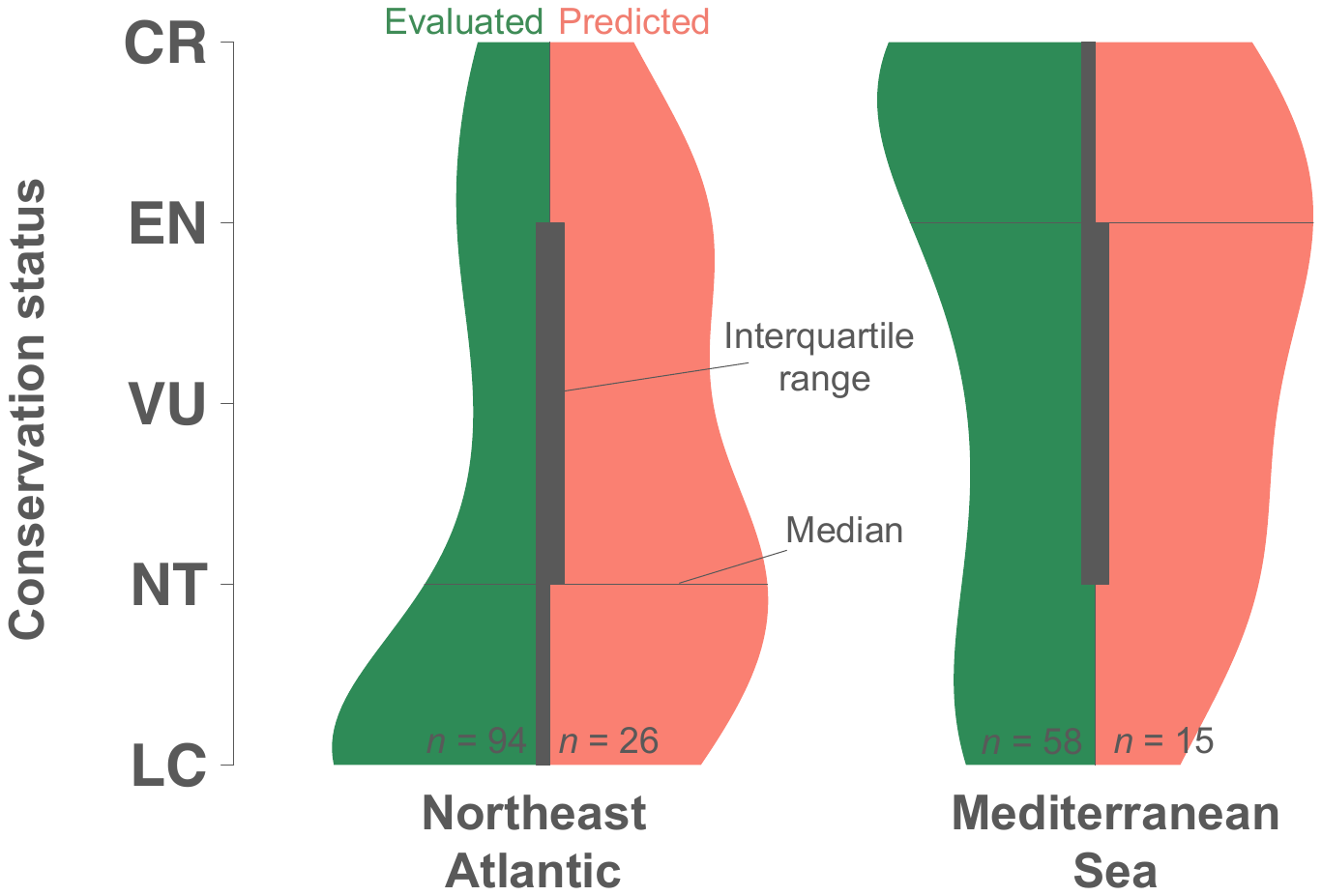

**Supplementary Figure A1**

**Distribution of evaluated and predicted shark and ray IUCN Red listings in Europe.** Split violin plots showing the distribution of Northeast Atlantic (left) and Mediterranean Sea (right) *evaluated* (left half) and *predicted* (right half) IUCN Red List categorisations of sharks and rays. There are 120 Northeast Atlantic shark and ray species, 94 of which were *evaluated* by the IUCN and 26 *predicted* for in this study. There are 73 Mediterranean Sea species, 58 of which were *evaluated* and 15 *predicted* for. The thick central vertical bars indicate the interquartile range of categorisations, while the horizontal lines show the median categorisation for each grouping. IUCN categories: CR = Critically Endangered, EN = Endangered, VU = Vulnerable, NT = Near Threatened, LC = Least Concern
